# Vagal Heart Rate Variability During Rapid Eye Movement Sleep Reduces Negative Memory Bias

**DOI:** 10.1101/2024.08.30.610388

**Authors:** Allison B. Morehouse, Katharine C. Simon, Pin-Chun Chen, Sara C. Mednick

**Author notes:** Correspondence should be addressed to: Sara C. Mednick.

## Abstract

Emotional memories change over time, but the mechanisms supporting this change are not well understood. Sleep has been identified as one mechanism that supports memory consolidation, with sleep selectively benefitting negative emotional consolidation at the expense of neutral memories, with specific oscillatory events linked to this process. In contrast, the consolidation of neutral and positive memories, compared to negative memories, has been associated with increased vagally-mediated vagal heart rate variability (HRV) during wakefulness. However, how HRV during sleep contributes to emotional memory consolidation remains unexplored. We investigated how sleep oscillations and vagal contributions during sleep contribute to the consolidation of neutral and negative memories. Using a double-blind, placebo-controlled, within-subject, cross-over design, we examined the impact of pharmacological vagal suppression using zolpidem on overnight emotional memory consolidation. Thirty-two participants encoded neutral and negative pictures in the morning, followed by picture recognition tests before and after a night of sleep. Zolpidem or placebo was administered in the evening before overnight sleep, and participants were monitored with electroencephalography and electrocardiography. In the placebo condition, greater overnight improvement for neutral pictures was associated with higher vagal HRV in both Non-Rapid Eye Movement Slow Wave Sleep (NREM SWS) and REM. Additionally, the emotional memory tradeoff (i.e., difference between consolidation of neutral versus negative memories) was associated with higher vagal HRV during REM, but in this case, neutral memories were remembered better than negative memories, indicating a potential role for REM vagal HRV in promoting a positive memory bias overnight. Zolpidem, on the other hand, reduced vagal HRV during SWS, increased NREM sigma power, and eliminated the positive memory bias. Lastly, we used a stepwise linear mixed effects regression to determine how NREM sigma power and vagal HRV during REM independently explained the variance in the emotional memory tradeoff effect. We found that the addition of vagal HRV in combination with sleep significantly improved the model’s fit. Overall, our results suggest that sleep brain oscillations and vagal signals synergistically interact in the overnight consolidation of emotional memories, with REM vagal HRV critically contributing to the positive memory bias.

## Introduction

The arc of healthy emotional memory processing changes over time, beginning with heightened emotional reactivity and intense reliving of the recent episodic experience, followed by the gradual reduction in affective response and concurrent strengthening of the memory for episodic details (van der Helm & Walker, 2011; Walker, 2010). Although the behavioral components of this process are somewhat understood, the underlying mechanisms driving this process are not. Several lines of evidence show that sleep may play important role by prioritizing the consolidation of negative memories at the expense of neutral memories, termed the emotional memory tradeoff effect (Nishida et al., 2009; van der Helm & Walker, 2011; Wagner et al., 2006; Wiesner et al., 2015). On the other hand, positive memories, at the expense of negative ones, have been shown to increase in association with autonomic activity during waking, specifically the vagal/parasympathetic component of heart rate variability (HRV) (Cho et al., 2023). However, no studies have investigated how HRV during sleep contributes to emotional memory processing. The current study investigates sleep specific neural and autonomic contributions to the consolidation of neutral and negative memories.

Although studies demonstrate the importance of sleep in emotional memory consolidation, there are mixed results as to the relative importance of the different sleep stages (Cox et al., 2018; Denis et al., 2022; Groch et al., 2013; Lipinska et al., 2019; Payne et al., 2008; Schäfer et al., 2020). Nap studies examining the differential effects of different sleep stages, have indicated that NREM is selectively related to the consolidation of negative, episodic information, with no added benefit from time spent in REM (Cellini et al., 2016; Payne et al., 2015). These findings align with overnight studies that pharmacologically enhanced specific NREM oscillatory activity, spindle activity (12-15 Hz), using zolpidem. They found that increased spindle activity was associated with greater memory for negative than neutral stimuli, compared with placebo (Kaestner et al., 2013; Simon et al., 2022).

However, other research emphasized a role of REM in emotional memory consolidation, with studies showing that REM-rich sleep, rather than SWS-rich sleep, enhances emotional text compared to neutral text (Ackermann & Rasch, 2014; Goldstein & Walker, 2014; Wagner et al., 2001). Further studies show positive correlations between negative emotional memory and minutes of REM during a daytime nap (Nishida et al., 2009) and overnight (Wiesner et al., 2015). Carr and Nielsen investigated the effect of REM on positive, negative, and neutral words and found that a nap with REM, compared to wake or a nap with only NREM, enhanced memory for positive, compared to negative, words (Carr & Nielsen, 2015). Similarly, another study demonstrated that a nap with REM, compared to wake, showed reduced ratings of fearful expressions and increased ratings of happy faces (Gujar et al., 2011). These results suggest that REM modulates emotional reactivity towards a positive bias. In contrast, REM deprivation, or decreasing the time spent in REM sleep, has led to decreased arousal ratings to negative stimuli, suggesting that REM may exacerbate negative emotional arousal (Lara-Carrasco et al., 2009). These mixed results on whether REM enhances positive or negative emotional stimuli have been reflected in meta-analyses, indicating that the effect of sleep and its stages may be context or content specific (Lipinska et al., 2019; Schäfer et al., 2020). Therefore, although NREM and REM have been associated with emotional memories, more research is needed to further explore this relationship and other potential mechanisms through which sleep processes emotional memories. We propose that these mixed results may in part stem from the lack of consideration of physiological mechanisms, with vagal HRV potentially helping to explain the discrepancies across studies.

The anticipated role of vagal HRV during sleep in emotional memory consolidation is supported by an extensive body of research that has investigated the relation between resting daytime HRV and emotion regulation (Malik & Camm, 1993; Sakaki et al., 2016). HRV, or variation in time between successive heartbeats, reflects the ability of the parasympathetic system to self-regulate in response to environmental change. Higher HRV signifies more effective potential to transition from a high sympathetic arousal state (e.g., during stressful situations) to a relaxed state (Malik & Camm, 1993). A recent study by Cho et al. (2023) examined how causally increasing resting HRV using HRV biofeedback affected the neural mechanisms of emotional memory processing (Cho et al., 2023). They determined that increasing HRV was associated with greater memory overall, as well as a memory bias favoring positive over negative images. This positive memory effect was mediated by changes in left amygdala-medial prefrontal cortex (mPFC) functional connectivity (Cho et al., 2023). Other studies that similarly increased HRV using a biofeedback intervention showed enhanced connectivity between the PFC and cortical regions involved in cognitive processes, such as executive functioning (Schumann et al., 2021). Indeed, studies have found an association between higher HRV and improved working memory (Hansen et al., 2003; Suriya-Prakash et al., 2015), enhanced decision making (Forte et al., 2021), efficient attentional control (Park et al., 2012), and better executive functioning (Thayer et al., 2009; Williams et al., 2019). Overall, these findings emphasize the role of HRV in emotion regulation and the enhancement of positive memories, which may be supported by broader executive functioning.

A growing body of research indicates that HRV during sleep also supports memory processes. Sleep strongly modulates ANS activity, with an overall reduction in heart rate and an increase in the parasympathetic component of HRV (high-frequency HRV: HF HRV; 0.15-0.4 Hz) during NREM (Chen et al., 2020b, 2020a; Trinder et al., 2012; Whitehurst et al., 2018). REM, in comparison, is a mixed autonomic state including both high sympathetic and parasympathetic activity (Chen et al., 2020; Chen et al., 2021). Critically, the autonomic activity during NREM and REM has been differentially related to improvement in sleep-dependent processing (Chen et al., 2020; Chen et al., 2021; Whitehurst et al., 2016). Whitehurst et al. (2016) examined the effect of implicit priming on enhancing post-nap performance on a creativity task and measured the sleep and HRV variables associated with performance improvement. They reported that HF HRV during REM and the total minutes spent in REM accounted for over 70 percent of the variance in overnight memory improvement (Whitehurst et al., 2016). Notably, REM minutes and HF HRV were not correlated with each other, suggesting that these features of sleep may be contributing non-overlapping mechanisms. Prior literature suggests a role for vagal HRV in offline cognitive enhancement, yet no studies have investigated the role of overnight vagal HRV in emotional memory processing.

The current study examined the impact of vagal activity, as measured by HF HRV, during sleep on emotional memory processing. In our study, participants were tested on an emotional memory task before and after a night of sleep that included placebo or pharmacological, zolpidem, intervention. As zolpidem results in HF HRV reduction during SWS, we were able to independently evaluate the contributions of sleep and HRV HRV activity on emotional memory consolidation. Given the role of HRV during wakefulness in the consolidation of neutral and positive memories, we hypothesized that higher vagal activity during sleep would improve memory for neutral images in the placebo condition. We hypothesized that reducing vagal activity in the zolpidem condition would increase the emotional memory bias, resulting in greater memory for emotional compared with neutral images. We further hypothesized that incorporating vagal activity during sleep into the existing model of NREM sigma power, as described by Simon et al., (2022), would significantly enhance the model’s accuracy in predicting the overnight consolidation of emotional memory.

## Methods

### Participants

Thirty-three healthy participants between the ages of 18 and 30 (*M*_age_ = 20.5 years, SD = 2.91, 16 females) provided informed consent, which was approved by the Western Institutional Review Board and the University of California, Riverside Human Research Review Board. All participants were healthy with no personal history of neurological, psychological, or other chronic illness and were naïve to or had limited contact with (< 2 lifetime use and no use in last year) zolpidem. Our exclusion criterion included irregular sleep/wake cycles, past history or present diagnosis of a sleep disorder, personal or familial history of diagnosed psychopathology, substance abuse/dependence, loss of consciousness greater than 2 minutes, a history of epilepsy, current use of psychotropic medications, non-correctable visual impairments, or any cardiac or respiratory illness that might affect cerebral metabolism. Participants were assessed for inclusion in-person using a modified Structured Clinical Interview (SCID) for the Diagnostic and Statistical Manual of Mental Disorders-IV (DSM-IV) (First et al., 1997), self-report questionnaires examining current and past health and wellness, and underwent a medical evaluation with the study physician, including a toxicology screening for schedule I and II drug substances. Participants were compensated monetarily for their participation.

### Experimental Design (see Fig. 1 for Schematic)

The current work utilized data previously published in two studies that addressed: 1) the effect of sigma activity on emotional memory consolidation (Simon et al., 2022), and 2) the effect of HRV on episodic and working memory (Chen, et al., 2021). In study 1, we showed that zolpidem, compared with placebo, increased sigma power and increased emotional memory tradeoff. In study 2, we showed that zolpidem, compared with placebo, suppressed HRV during SWS and increased a tradeoff between episodic and working memory. The present study was designed to examine the unique effect of suppressing vagal activity during sleep on overnight emotional memory consolidation. We used a double-blind, placebo-controlled, within-subject, cross-over design, with visit order randomized and counterbalanced, to examine the role of HRV during sleep on sleep-dependent emotional memory processing.

### Procedures

All participants reported to the lab between 8:00 AM and 9:00 AM and completed the encoding phase of the Emotion Picture Task (EPT). Afterwards, participants remained in the lab until 2:00 PM to undergo hourly monitoring of their vitals (blood pressure, heart rate, and subjective measurements). Participants were asked to refrain from exercise, sleep, caffeine, or other drug substances between sessions. Participants returned to the lab at 9:00 PM to complete EPT Test 1. Participants were then given 10-mg dose of placebo or zolpidem and polysomnography was administered. Vitals were monitored again until lights out at approximately 11:00 PM. All participants were woken up by 9:00 AM and provided breakfast and a break. EPT Test 2 was administered at approximately 10:30 AM. Participants completed this protocol twice, once for each drug condition (see Fig. 1 for schematic).

### Drug Protocol

Directly before lights out, participants ingested either 10-mg of placebo or zolpidem, a positive allosteric modulator of GABAA receptors with a short half-life (1.5–4.5 h) and rapid onset which was prepared by the MDMX Corona Research Pharmacy. Zolpidem is a GABAA agonist that has been shown to decrease average heart rate (Sinha & Yadav, 2020) and overall HRV (Machado et al., 2011, Chen et al., 2021), while increasing relative fast sigma power and sleep spindle density (Mednick et al., 2013; Simon et al., 2022; Zhang et al., 2020). The zolpidem and the placebo capsules were visually indistinguishable.

### Polysomnography

This study used a 32-channel electroencephalography (EEG) cap (EASYCAP GmbH) with Ag/AgCI electrodes placed according to the international 10–20 System (Jasper, 1958). There were 22 neural channels, 2 electrocardiogram (ECG), 2 electromyogram (EMG), 2 electrooculogram (EOG), a ground, a reference channel, and 2 mastoid recordings. EEG was recorded at a 1000-Hz sampling rate and preprocessed in BrainVision Analyzer 2.0 (BrainProducts, Munich Germany). During preprocessing, all neural electrodes were re-reference to the contralateral mastoid and artifacts and arousals were removed via visual inspection. Low pass (0.3 Hz) and high-pass filters (35 Hz) were applied to all neural and EOG channels. The raw data was scored using eight neural electrodes (F3, F4, C3, C4, P3, P4, O1, O2), EMG, and EOG according to Rechtschaffen and Kales in 30-s epochs (Rechtschaffen & Kales, 1968). Then, wake after sleep onset (WASO) and minutes in each sleep stage (NREM Stage 1, NREM Stage 2, NREM Stage 3, and REM) were calculated. Simon et al. (2022) examined the impact of sleep frequency bands on overnight emotional memory consolidation. Following their protocol, we analyzed relative fast sigma power (12-15 Hz) from the central channels (C3 and C4) in NREM N2 and SWS, given that sigma power is most prominent over the central channels (Simon et al., 2022). Based on their results, we focused on C4 in these set of analyses.

### Emotion Picture Task (EPT)

#### Stimuli

The EPT consisted of three phases: an encoding phase and two retrieval phases, see Figure 1. The pre-sleep retrieval phase was in the evening, approximately 12-hours after encoding (Test 1), and the post-sleep retrieval phase was the next morning after a night of sleep in the lab and 24-hours after encoding (Test 2). Participants were shown and asked to remember negative and neutral pictures from the International Affective Picture System, or IAPS (Lang et al., 1997). The standard IAPS ratings ranged from 1, more unpleasant and less arousing, to 9, more pleasant and highly arousing. These standard IAPS ratings were used to control for arousal and valence to maximize the difference between neutral, low arousing (valence: *M* = 5.13, *SD* = 1.44; arousal: *M* = 3.87, *SD* = 2.19) and negative, highly arousing (valence: *M* = 2.88, *SD* = 1.57; arousal: *M* = 5.52, *SD* = 2.09) pictures. The negative and neutral pictures were counterbalanced across condition without repetition. This task was administered on a Windows computer using Matlab with Psychtoolbox. Our task code (encoding and recognition test) is freely available online at https://github.com/MednickLab/GenericMemoryTask.

**Figure 1.**
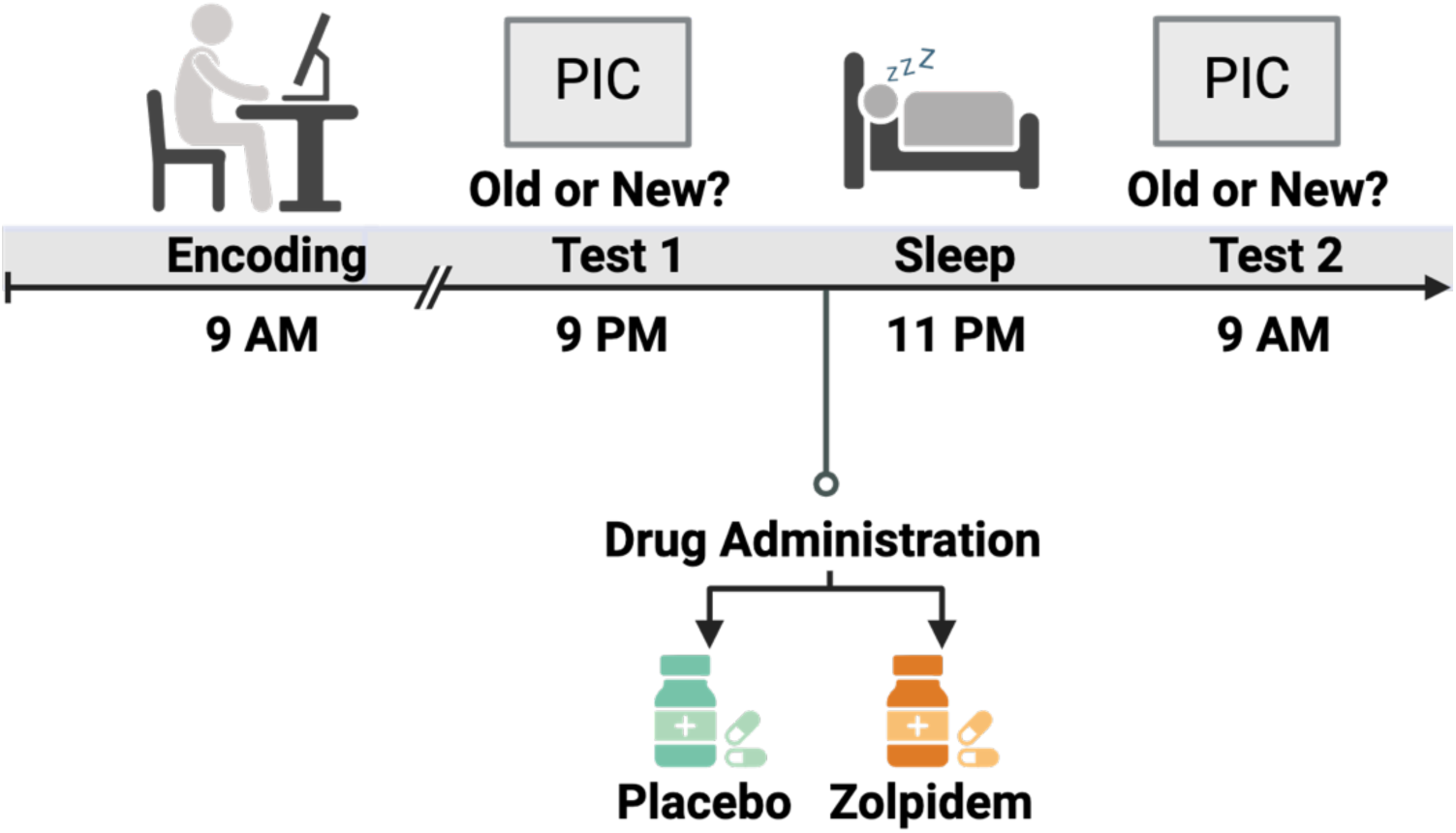
Experimental Design Schematic. This figure was created using BioRender.com

**Figure 1.**
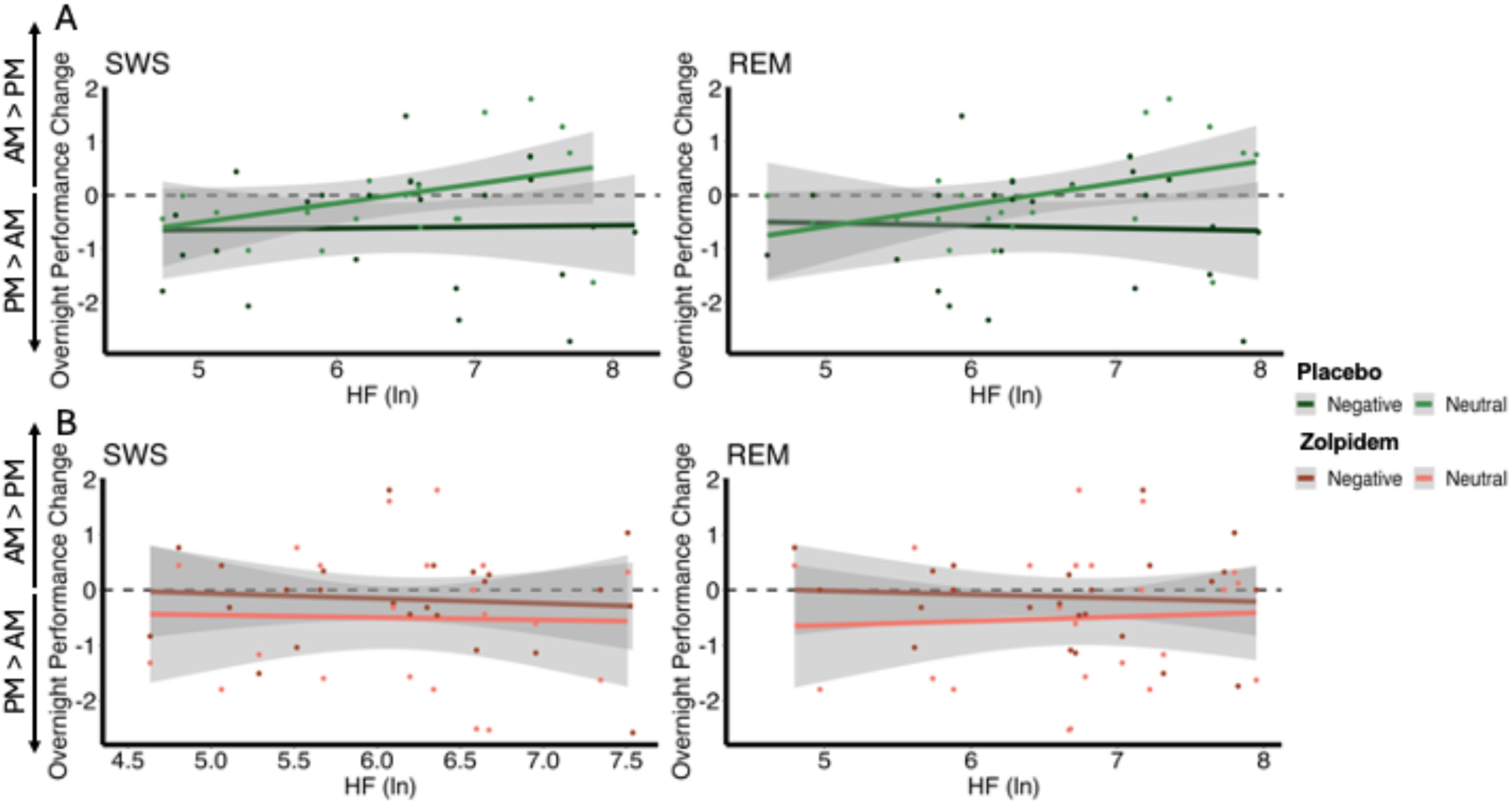
Correlations Between Vagal Activity and the Overnight Performance Change in Emotional and Neutral Memory. (A) In the placebo condition, overnight performance change for neutral images positively correlated with HRV during SWS (r=0.520, p_adj_=.011) and with HRV during REM (r=0.550, p_adj_=.010). However, no significant associations were found between negative images and HRV during SWS (r=-0.057, p_adj_=1) or with HRV during REM (r=-0.099, p_adj_=1). (B) In the zolpidem condition, there were no significant correlations between overnight performance change for neutral or negative images and HRV during SWS (all p_adj_=1) or with HRV during REM (all p_adj_=1).

#### Encoding and Testing Procedure

We controlled for primacy and recency effects by displaying 4 neutral pictures each before and after the stimulus set. During the encoding phase, participants were shown 20 negative and 20 neutral pictures for 500ms. A fixation marker was always presented for 1000ms before each picture. Each retrieval test involved half of the initially presented pictures along with 20 new pictures – 10 negative and 10 neutral. For each picture, participants had unlimited time to determine whether it was old (shown at encoding) or new. See Figure 2 for EPT task example.

**Figure 2.**
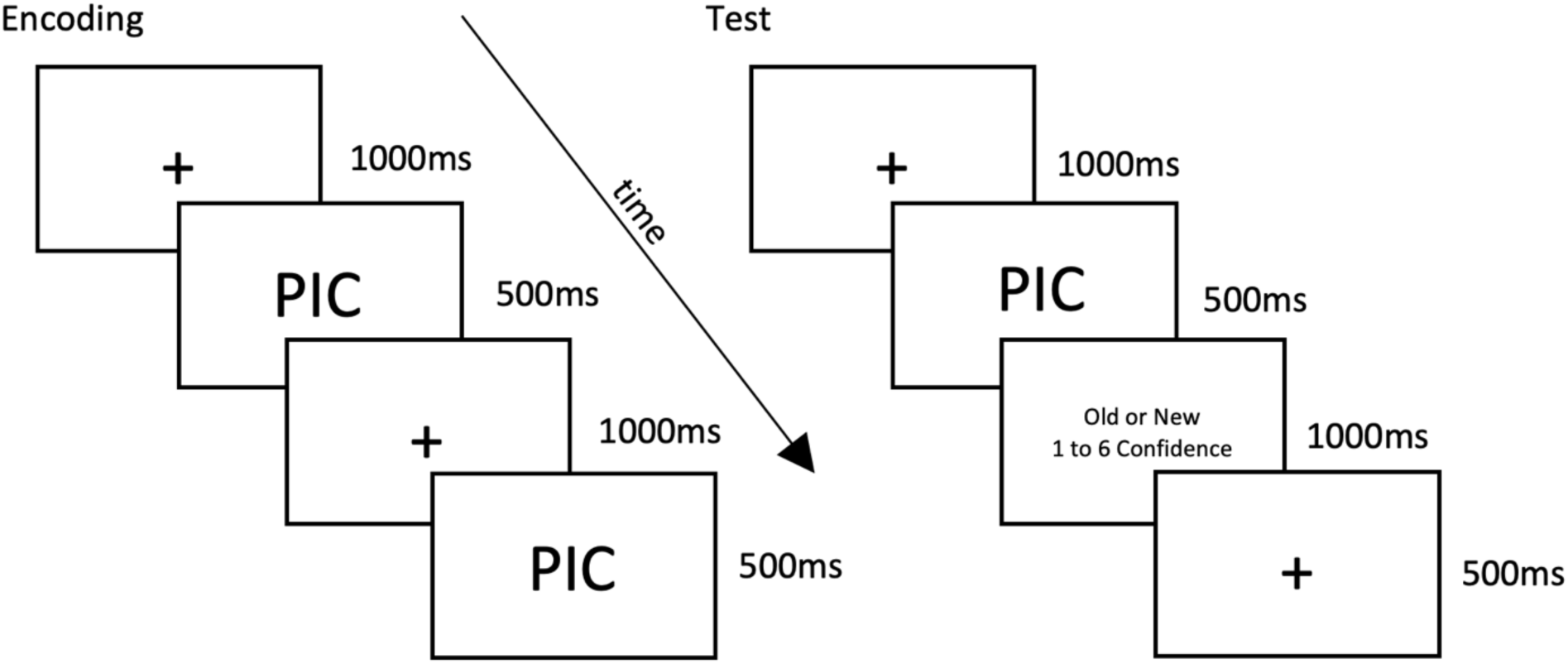
Emotional Picture Task Paradigm

**Figure 2.**
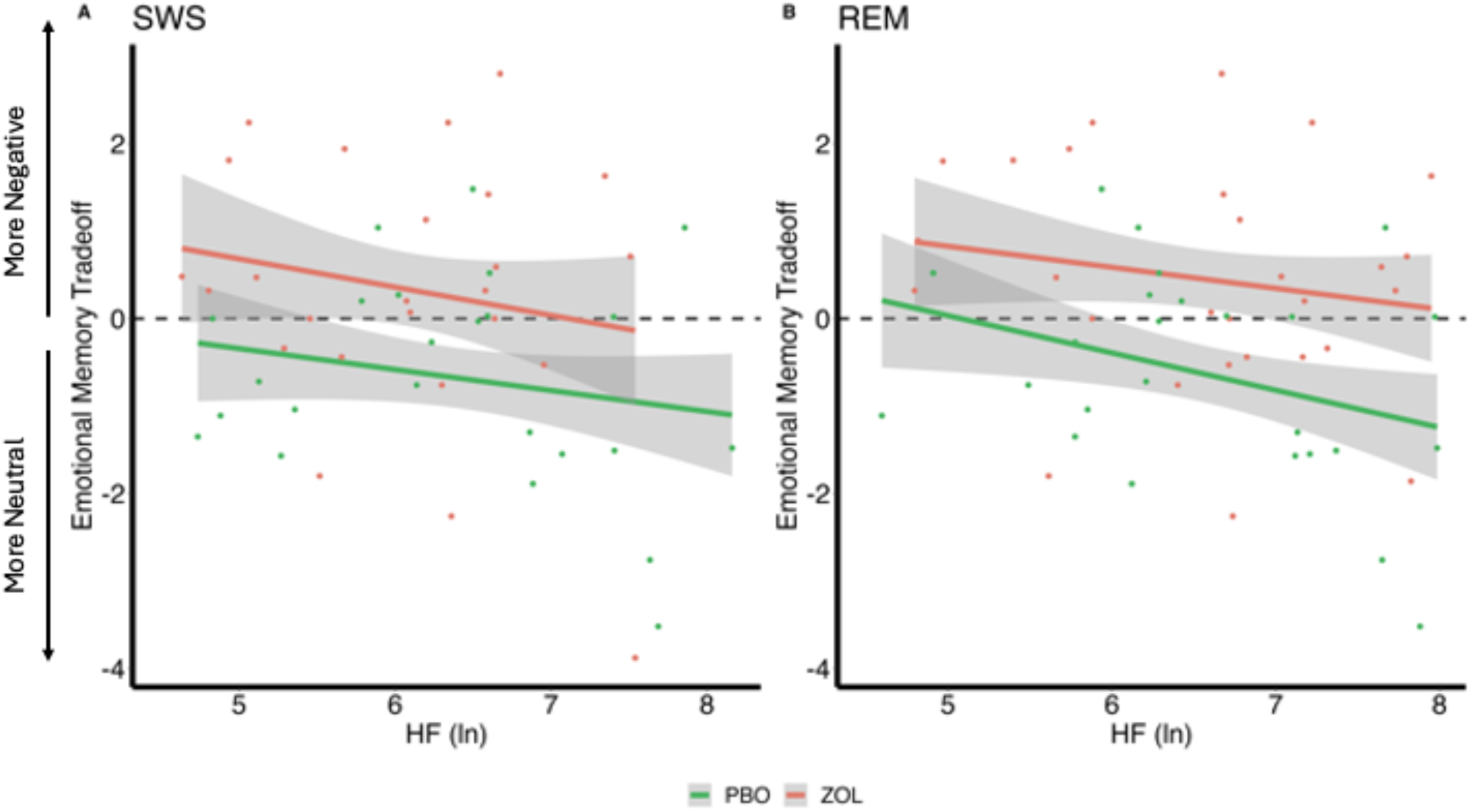
Correlations Between Vagal Activity and the Emotional Memory Tradeoff Effect. (A) There were no significant correlations between the emotional memory tradeoff effect and HRV during SWS for either the placebo (r=-0.36, p_adj_=.069) or zolpidem condition (r=-0.26, p_adj_=.38). (B) In the placebo condition, the emotional memory tradeoff effect negatively correlated with HRV during REM (r=-0.46, p_adj_=.034), but this association was eliminated by the zolpidem (r=-0.17, p_adj_=.38).

#### Performance Measures

Memory performance using discriminability index (d’) was calculated using the z transform of hit rate (% of pictures correctly identified as present) minus the false alarm rate (% of pictures incorrectly identified as present) at both Test 1 and Test 2 (see Supplementary Materials). Higher d’ scores signify better performance. We calculated overnight memory performance by the difference between Test 1 and Test 2 d’ (Test 2 d’ – Test 1 d’) separately for neutral and negative pictures. More positive overnight memory performance indicates better performance for the morning test compared to the evening test. We also created an emotional memory tradeoff score derived by subtracting the neutral overnight memory performance from the negative overnight memory performance (negative d’ difference– neutral d’ difference). For this score, positive scores would suggest a negative memory bias (negative better than neutral), whereas negative scores would suggest positive memory bias (neutral better than negative).

### Heart Rate Variability (HRV)

ECG data was collected using a modified Lead II Einthoven configuration. We analyzed HRV of the R-waves series during the night using Kubios HRV Analysis Software 2.2 (Biosignal Analysis and Medical Imaging Group, University of Kuopio, Finland), according to the Task Force guidelines (Electrophysiology Task Force of the European Society of Cardiology and the North American Society of Pacing and, 1996). The Kubios software automatically detected R-peaks of the QRS signal on the ECG. The identified R-peaks were visually examined, and incorrect R-peaks were manually edited by trained technicians. Artifacts were removed by the Kubios software using the provided automatic medium filter. Missing beats were corrected via cubic spline interpolation and then inter-beat intervals were calculated. Additionally, a third-order polynomial filter was applied to the time series to remove trend components. The RR intervals (ms; time interval between consecutive R-peaks) were analyzed with a MATLAB-based algorithm (Chen et al., 2021; Whitehurst et al., 2018) using an autoregressive model, with the model order set at 16 (Boardman et al., 2002).

#### HRV Measures

We calculated and selected HF HRV (0.15–0.40 Hz; ms^2^) as the overall measure of HRV because it is considered an index of vagal tone, and previous research has shown that zolpidem selectively suppresses vagal (HF HRV) tone, but has no impact on low-frequency (LF; 0.04 to 0.15 Hz; ms^2^) HRV (Chen et al., 2021; Machado et al., 2011). HF had skewed distributions, and therefore, it was transformed by taking the natural logarithm. HRV was calculated based on consecutive artifact-free 5-minute windows to remain consistent with previous nocturnal sleep studies (Ako et al., 2003; Brandenberger et al., 2003; de Zambotti et al., 2014). The entire 5-minute window needed to be free from stage transitions, arousal, or movements. We calculated and analyzed HRV for NREM SWS and REM.

### Data Reduction and Cleaning

The initial dataset included 33 records for each drug condition. Two participants withdrew from the study before completing both drug conditions due to scheduling conflicts – one did not complete the zolpidem visit and the other did not complete the placebo visit. Whenever possible, we retained the data for the visit these two participants did complete. Therefore, 31 participants successfully completed the entire study, including both drug conditions.

#### EPT

Negative or neutral performance data was removed when there were fewer than five out of ten hits, to exclude participants that performed below chance. Therefore, negative performance data was removed for two participants in the placebo condition and three participants in the zolpidem condition. Neutral performance data was removed for four participants in the placebo condition and four participants in the zolpidem condition. After cleaning, the placebo condition contained 30 participants’ negative performance data and 28 participants’ neutral performance data. After cleaning, the zolpidem condition contained 29 participants’ negative performance data and 28 participants’ neutral performance data. For between drug condition analyses, only the 31 participants that completed both visits were used: 23 participants after cleaning.

#### HRV

HRV data was lost due to experimental error for both visits for two participants and for the placebo visit of another two participants. The autoregressive model used to calculate HRV metrics required consecutive undisturbed windows of sleep without stage transitions, arousals, or movement, which resulted in some participants lacking sufficient HRV data for a sleep stage. For SWS sleep, the dataset included HRV data for 28 participants in the zolpidem condition and 27 in the placebo group before cleaning. For REM, the dataset included HRV data for 29 participants in the zolpidem condition and 27 in the placebo group before cleaning. We cleaned the data for each drug condition independently. We removed HRV data that were 2.5 standard deviations above or below the mean for that sleep stage before we took the natural logarithm. This resulted in the exclusion of one participant’s SWS HRV in the placebo condition, one participants’ SWS HRV in the zolpidem condition, and one participants’ REM HRV in the zolpidem condition. After data cleaning, the final dataset included SWS HRV data for 27 participants in the zolpidem condition and 26 in the placebo group and REM HRV data for 28 participants in the zolpidem condition and 27 in the placebo group.

### Statistical Analyses

All analyses were performed in R version 4.3.2 using packages lme4, ggplot2, dplyr, tidyr, lme4, and rstatix.

#### Associations between Vagal Activity and Emotional and Neutral Memory

We first computed Pearson’s *r* to assess the relation between HRV within each sleep stage and overnight memory performance, both for negative and neutral images. We also calculated the 95% confidence interval for the correlation and the t-statistic, both of which test the null hypothesis that there is no significant correlation between the two variables. These relations were assessed separately for each drug condition (zolpidem and placebo). In addition, we statistically compared the correlations between HRV within each sleep stage and negative versus neutral memory performance using the Fisher r-to-z transformation. We computed the emotional difference score (EmoDiff) by subtracting the fisher-transformed z-scores for negative and neutral stimuli (negative z-score – neutral z-score). The magnitude of the EmoDiff score indicates the size of the difference in standard deviation units. A positive EmoDiff score indicates that the correlation between HRV and memory performance for negative images is stronger than the correlation between HRV and memory performance for neutral images. A negative EmoDiff score indicates that the correlation between HRV and memory performance for neutral images is stronger than the correlation between HRV and memory performance for negative images. We corrected for multiple comparisons using the Holm-Bonferroni method, corrected correlations were statistically significant if p_adj_<0.05.

#### Vagal Activity and the Emotional Memory Tradeoff Effect

We computed Pearson’s *r*, a 95% confidence interval, and a test t-statistic to assess the relation between HRV within each sleep stage and emotional memory tradeoff score. These relations were assessed separately for each drug condition (zolpidem and placebo). We corrected for multiple comparisons using the Holm-Bonferroni method, corrected correlations were statistically significant if p_adj_<0.05.

#### Comparing the Impact of Sigma Power and Vagal Activity on the Emotional Memory Tradeoff Effect

Building off the findings of Simon et al., we used a regression framework to determine whether including HRV and sigma power in the model explains a significant amount of variance in the emotional memory tradeoff effect. We ran stepwise linear mixed effects models, first with a base model that included the emotional memory tradeoff as the dependent variable, condition and mean centered weight as covariates, and a random fixed effect for each subject. We included mean centered weight as a covariate in each of these analyses to account for the different rates of drug absorption due to the weight of our participants (Simon et al., 2022). The sigma model included SWS sigma power and an interaction term between condition and sigma power. The HRV model expanded upon the sigma model by including REM HRV and another interaction term between condition and REM HRV. The interaction terms allowed us to determine whether the effect of condition on the emotional memory tradeoff effect varies depending on REM HRV or sigma power. For model selection, we calculated each model’s adjusted marginal R^2^. Since all models include a common random effect term for each subject, we used the adjusted marginal R^2^, rather than the conditional R^2^, because it represents the portion of variance explained by the fixed effects only and is more in line with the intuitive definition of R^2^ (Sotirchos et al., 2019). We then ran likelihood ratio tests between the sigma model and the full model that included both sigma power and HRV.

## Results

We first confirmed the HRV results as published in Chen et al. with zolpidem significantly reducing HRV during SWS (Chen et al., 2021). Second, in the placebo condition, we confirmed the behavioral results of greater overnight consolidation of neutral memories than negative memories (Simon et al., 2022). Third, in the zolpidem condition, we replicated the result that there was greater overnight retention of negative memories compared to neutral memories (Simon et al., 2022). For the replications of prior literature, see the supplementary materials.

**Table 1.** Summary of Sigma and HRV Parameters Across Sleep Stages.

|  | N2 |  |  |  |  | SWS |  |  |  |  | REM |  |  |  |  |
| --- | --- | --- | --- | --- | --- | --- | --- | --- | --- | --- | --- | --- | --- | --- | --- |
|  | PBO |  | ZOL |  |  | PBO |  | ZOL |  |  | PBO |  | ZOL |  |  |
| Measure | Mean | SD | Mean | SD | p | Mean | SD | Mean | SD | p | Mean | SD | Mean | SD | p |
| C4 Spindle Power | 1.423 | 0.551 | 1.722 | 0.749 | **0.006** | 0.468 | 0.214 | 0.631 | 0.398 | **0.002** | - | - | - | - | - |
| C3 Spindle Power | 1.417 | 0.534 | 1.766 | 0.801 | **0.002** | 0.525 | 0.257 | 0.653 | 0.437 | **0.042** | - | - | - | - | - |
| HF (ln) | 6.568 | 0.945 | 6.487 | 0.935 | 0.545 | 6.058 | 1.105 | 6.060 | 1.094 | **0.034** | 6.445 | 1.011 | 6.446 | 1.001 | 0.133 |

### Associations between Vagal Activity and Emotional and Neutral Memory

Here, we examined overnight performance change for negative and neutral pictures in relation to HRV during SWS and REM in both the placebo and zolpidem conditions, using Pearson’s *r* correlations.

#### Placebo

Overnight performance change (Test 2 – Test 1) for neutral pictures was positively correlated with SWS HRV (r=0.520, p_adj_=.011) and REM HRV (r=0.550, p_adj_=.010). Individuals with higher vagal HRV during SWS and REM showed better memory performance for neutral pictures in the morning, given their pre-sleep baseline performance, compared to individuals with lower vagal autonomic tone. We found no significant results for HRV and the overnight performance change for negative pictures (all p_adj_=1), suggesting a selective enhancement of neutral memories over negative with increased HRV during SWS and REM.

#### Zolpidem

As predicted, the reduction in HRV in the zolpidem condition eliminated the bias towards neutral memories that was found in the placebo condition. As such, we found no significant correlations between the overnight performance change for neutral images and HRV in any sleep stage (all p_adj_=1). There were also no significant correlations between the overnight performance change for negative images and HRV in any sleep stage (all p_adj_=1). This suggests that significant reductions in HRV impairs the overnight processes that lead to a bias towards neutral memories.

**Table 2.** Correlations Between Vagal Activity and the Emotional and Neutral Memory.

|  |  |  | SWS |  |  |  | REM |  |  |  |
| --- | --- | --- | --- | --- | --- | --- | --- | --- | --- | --- |
| EPT | HRV | Drug | t | r | p<sub>adj</sub> | CI | t | r | p<sub>adj</sub> | CI |
| Neutral Overnight Performance Change | HF (ln) | PBO | 2.778 | 0.520 | **0.011** | [0.135, 0.766] | 3.089 | 0.550 | **0.010** | [0.188, 0.780] |
| Neutral Overnight Performance Change | HF (ln) | ZOL | 0.525 | 0.110 | 1.000 | [-0.306, 0.492] | 0.292 | 0.060 | 1.000 | [-0.336, 0.436] |
| Negative Overnight Performance Change | HF (ln) | PBO | -0.273 | -0.057 | 1.000 | [-0.442, 0.346] | -0.478 | -0.099 | 1.000 | [-0.476, 0.308] |
| Negative Overnight Performance Change | HF (ln) | ZOL | -0.427 | -0.089 | 1.000 | [-0.468, 0.317] | -0.348 | -0.072 | 1.000 | [-0.455, 0.332] |

To further investigate this potential positive memory bias with higher vagal autonomic function, we tested whether the correlations for memory performance and HRV measures were significantly different between negative and neutral memories. We computed the EmoDiff score, or the Holm–Bonferroni corrected Fisher r-to-z transformations, to compare the differences between these correlations.

#### Placebo

We found significant differences between the correlations between HRV and negative overnight performance versus neutral overnight performance change, for both SWS (EmoDiff = - 2.240, p_adj_=.025) and REM (EmoDiff = −2.538, p_adj_=.022). These results suggest that higher HRV across sleep stages predicted a selective enhancement in overnight memory for neutral compared with negative memories, indicative of a reduced negative memory bias, or an increased positive memory bias.

#### Zolpidem

Again, zolpidem, which significantly lowered vagal autonomic tone, eliminated the positive memory bias for neutral over negative memories. For the zolpidem condition, there were no significant differences in the relation between HRV and the overnight performance change for neutral or negative images (all p_adj_=.926).

**Table 3.** Differences between Fisher-Transformed Z-scores for Vagal Activity and Emotional and Neutral Memory.

|  |  |  | SWS |  | REM |  |
| --- | --- | --- | --- | --- | --- | --- |
| EPT | HRV | Drug | p<sub>adj</sub> | EmoDiff | p<sub>adj</sub> | EmoDiff |
| Overnight Memory Consolidation | HF (ln) | Pbo | 0.025 | -2.240 | 0.022 | -2.538 |
| Overnight Memory Consolidation | HF (ln) | Zol | 0.926 | -0.734 | 0.926 | -0.486 |

### Vagal Activity and the Emotional Memory Tradeoff Effect

Given that we demonstrated a unique contribution of HRV during sleep to negative and neutral memory processing, we, next, examined the emotional memory tradeoff score (negative overnight performance change – neutral overnight performance change) to investigate a possible trade-off between higher HRV and greater neutral memory at the expense of negative memory. As explained in the Methods, more positive scores indicate a negative memory bias (negative better than neutral), whereas negative scores indicate a positive memory bias (neutral better than negative).

#### Placebo

There was a significant negative correlation between the emotional memory tradeoff score and REM HRV (r=-0.460, p_adj_=.034), that is, individuals with higher HRV during REM exhibited greater memory for neutral images at the expense of negative images (positive overnight memory bias). No significant correlation for SWS HRV (r=-0.360, p_adj_=.069) was found.

#### Zolpidem

Adding zolpidem eliminated significant associations between the emotional memory tradeoff score and HRV measures during SWS (r=-0.260, p_adj_=.380) and REM (r=-0.170, p_adj_=.380). These findings suggest that reducing HRV neutralizes the positive memory bias, leading to a non-significant negative correlation where negative images are remembered more.

**Table 4.**
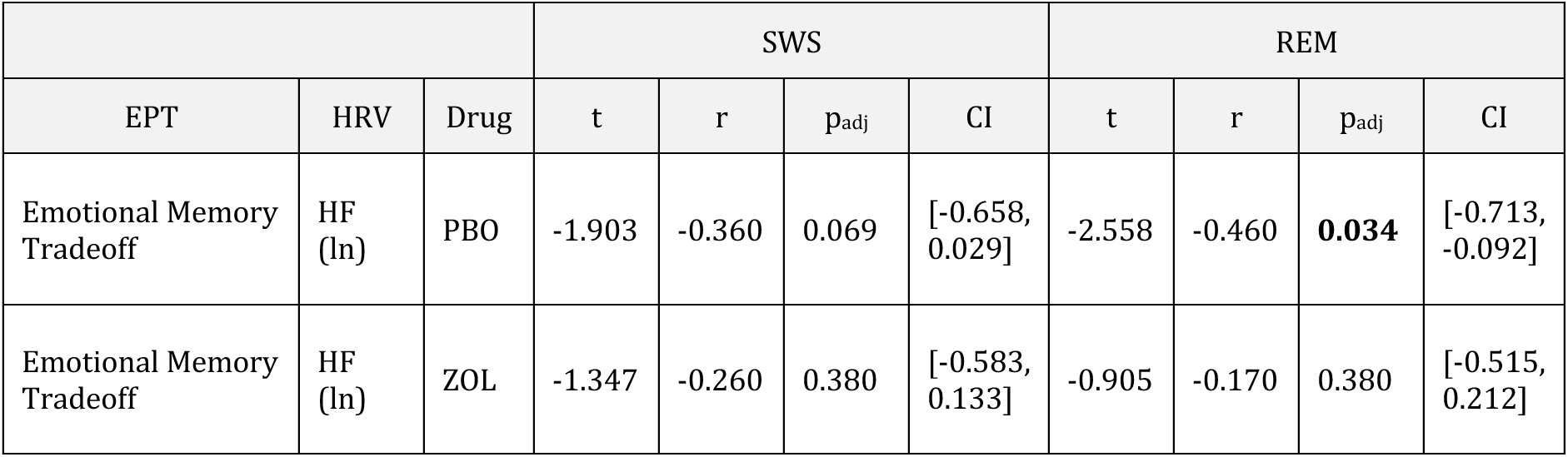
Correlations Between Vagal Activity and the Emotional Memory Tradeoff Effect.

|  |  |  | SWS |  |  |  | REM |  |  |  |
| --- | --- | --- | --- | --- | --- | --- | --- | --- | --- | --- |
| EPT | HRV | Drug | t | r | p<sub>adj</sub> | CI | t | r | p<sub>adj</sub> | CI |
| Emotional Memory Tradeoff | HF (ln) | PBO | -1.903 | -0.360 | 0.069 | [-0.658, 0.029] | -2.558 | -0.460 | **0.034** | [-0.713, -0.092] |
| Emotional Memory Tradeoff | HF (ln) | ZOL | -1.347 | -0.260 | 0.380 | [-0.583, 0.133] | -0.905 | -0.170 | 0.380 | [-0.515, 0.212] |

### Comparing the Impact of Sleep Spindles and Vagal Activity on the Emotional Memory Tradeoff Effect

Given the concurrent enhancing effects on spindles by zolpidem, we examined the degree to which both spindles and HRV contribute to the emotional memory trade-off effect using a regression framework. The base model tested the effect of drug condition, the sigma model tested the effect of SWS sigma power and its interaction with condition, and the full model added REM HRV and its interaction with condition. Using likelihood ratio tests, we determined whether the addition of REM HRV significantly improved the model’s explanatory power. This analysis helped quantify the additional variance in the emotional memory tradeoff effect accounted for by HRV, offering deeper insights into its physiological influences.

#### Base Model

We ran stepwise linear mixed effects models with a base model including the emotional memory tradeoff as the dependent variable, drug condition and centered weight as covariates, and a random fixed effect for each subject. The base model determined the effect of drug condition and centered weight on the emotional memory tradeoff effect. For the base model, drug condition had a statistically significant effect on emotional memory tradeoff (β = 0.853, t = 2.731, p = .010). This result indicates that the zolpidem conditions, compared to placebo, had a higher emotional memory tradeoff score, and therefore, greater memory for negative pictures at the expense of neutral pictures. There was not a significant effect of centered weight (β < 0, t = −0.053, p = .958). The base model had an adjusted marginal R^2^ of 0.077.

#### Sigma Model

Next, we included SWS sigma power and an interaction between condition and sigma power into the base model. There was a significant interaction between condition and sigma power during SWS (β = −2.618, t = −2.110, p = .042). Thus, as previously shown, the impact of sigma power on the emotional memory tradeoff score significantly differed between the zolpidem and placebo conditions, with stronger correlation in zolpidem than placebo (negative β coefficient). Additionally, there was a significant fixed effect for drug condition (β = 2.164, t = 3.148, p = .004), indicating that, compared to placebo, zolpidem had greater emotional memory tradeoff towards negative pictures. There was not a significant fixed effect for sigma power (β = 2.159, t = 1.697, p = .096). The fast sigma model had an adjusted marginal R^2^ of 0.127.

#### Full Model

Here, we tested the emotional memory tradeoff score as the dependent variable and drug condition, centered weight, SWS sigma power and REM HRV as covariates, and a random effect of subject ID. This model also included two interactions terms: an interaction between condition and SWS sigma power and an interaction between condition and REM HRV. After controlling for the separate effects of condition and centered weight, there was a significant fixed effect for REM HRV (β = −0.563, t = −2.753, p = .008) and for sigma power (β = 3.322, t = 3.192, p = .003). The significant interaction between condition and sleep sigma power during SWS remained (β = −12.837, t = −2.387, p = .021). Yet, there was not a significant interaction between condition and REM HRV (β = 0.253, t = 0.797, p = .429) or a significant fixed effect for drug condition (β = 0.693, t = 0.330, p = .743). These results suggest that while REM HRV and SWS sigma power independently contribute to the emotional memory tradeoff, the interaction between REM HRV and condition does not significantly influence memory consolidation. This indicates that REM HRV has a distinct role in emotional memory consolidation and its effect was not significantly modulated by the drug condition. This model had an adjusted marginal R^2^ of 0.365.

The sigma model built upon the base model to also include SWS sigma power and explained approximately 12.7% of the variance based on the fixed effects. By adding REM HRV, the full model explained approximately 36.5% of the variance. We ran a likelihood ratio test to compare the sigma and full models and showed that including HRV in the model significantly improves its fit (χ² =9.4603, p = .008), providing evidence that HRV contributes to the explanation of variance in the emotional memory tradeoff effect. Therefore, REM HRV is a crucial predictor for overnight emotional memory consolidation and should be included in models aiming to understand this process.

**Table 5.** Model Comparison Results. Bold text indicates significance at *p* ≤ .05. We used adjusted marginal R², reflecting variance explained by fixed effects, as the means of model comparison. Overall, the full model including both REM HRV and SWS sigma was the best fit for the data and significantly improved the fit (χ² =9.4603, p = .008).

| Model | Term | Estimate | Std. Error | t | p | Marginal R<sup>2</sup> |
| --- | --- | --- | --- | --- | --- | --- |
| Base Model | Intercept | -0.660 | 0.268 | -2.451 | **0.017** | 0.076 |
|  | Condition<sub>ZOL</sub> | 0.852 | 0.312 | 2.730 | **0.010** |  |
|  | Centered Weight | -0.001 | 0.007 | -0.053 | 0.958 |  |
| Sigma Model | Intercept | -1.613 | 0.656 | -2.457 | **0.017** | 0.126 |
|  | Condition<sub>ZOL</sub> | 2.164 | 0.687 | 3.147 | **0.003** |  |
|  | SWS C4 Sigma | 2.159 | 1.272 | 1.696 | 0.095 |  |
|  | Centered Weight | 0.001 | 0.007 | 0.141 | 0.888 |  |
|  | Condition * SWS C4 Sigma | -2.618 | 1.240 | -2.111 | **0.042** |  |
| Full Model: HRV & Sigma Power | Intercept | 1.587 | 1.392 | 1.140 | 0.260 | 0.365 |
|  | Condition<sub>ZOL</sub> | 0.692 | 2.098 | 0.330 | 0.742 |  |
|  | SWS C4 Sigma | 3.322 | 1.040 | 3.191 | **0.002** |  |
|  | REM HF (ln) | -0.564 | 0.204 | -2.754 | **0.008** |  |
|  | Centered Weight | 0.005 | 0.005 | 1.068 | 0.290 |  |
|  | Condition * SWS C4 Sigma | -2.837 | 1.188 | -2.387 | **0.021** |  |
|  | Condition *REM HF (ln) | 0.253 | 0.317 | 0.797 | 0.429 |  |

## Discussion

We investigated the impact of vagal activity during sleep on overnight memory consolidation for emotional and neutral pictures. To achieve our goal, we tested participants on an emotional picture task before and after a night of sleep where we measured both brain oscillatory activity and vagal HRV. Employing a within subjects, double blind, crossover design, we assessed the causal role of HRV through a pharmacological intervention (zolpidem vs placebo) that decreased vagal HRV during slow wave sleep (SWS) (Chen et al., 2021). Building on prior findings (Simon et al., 2022), we found that the placebo condition had enhanced neutral and reduced negative memories, while the zolpidem condition showed the reverse effect, enhanced negative compared to neutral memories. Second, in the placebo condition, we found that the overnight neutral memory improvement was positively associated with vagal HRV during SWS and REM, but did not find a similar association between HRV and negative memories. Pharmacologically suppressing overnight vagal activity with zolpidem eliminated associations between vagal HRV during sleep and neutral memory consolidation and shifted overnight memory consolidation toward a negativity bias. Third, we examined associations between HRV during SWS and REM and the emotional memory tradeoff measure for each drug condition. In the placebo condition, higher REM HRV, but not SWS HRV, was significantly correlated with greater memory for neutral images at the expense of negative images, whereas the association was absent in the zolpidem condition. Finally, using a stepwise linear regression model, we found that REM HRV and NREM sigma power explained independent variance in the emotional memory tradeoff effect. Together, our results implicate a mechanistic role for vagal HRV during REM in shifting overnight memory consolidation away from negative experiences.

The expected emotional memory tradeoff, where emotional memories are remembered better than neutral (Nishida et al., 2009; Wagner et al., 2001, 2006; Wiesner et al., 2015), was not shown in our placebo condition (see supplementary materials, Simon et al., 2022). Similar null or reversed findings (neutral better than negative) have been reported in other research (Cellini et al., 2016; Kaestner et al., 2013; Lipinska et al., 2019; Simon et al., 2022). Meta-analyses also report inconsistent results and suggest that the variability in findings may be attributable to differing study design (e.g., free recall versus recognition, sleep and wake control conditions, etc.) and sample characteristics (i.e., age, sex) (Lipinska et al., 2019; Schäfer et al., 2020).

Importantly, the expected tradeoff emerged in our pharmacological intervention, which both reduced vagal HRV and increased NREM spindle activity; and our regression models demonstrated that both REM HRV and spindle activity explained significant and independent variance in the drug conditions by memory tradeoff interaction. Kaestner et al. (2013) found similar results with zolpidem-driven increases in sigma activity associated with greater negative than neutral memories. Given the fact that HRV was not measured in prior studies, the current results suggest an intriguing possibility that autonomic activity may be a heretofore unexamined third variable that could partially explain discrepancies across studies.

Our findings that HRV during sleep modulated the emotional memory tradeoff align with research on the role of resting HRV (assessed during wake) for emotional memory and self-regulation (Cho et al., 2023; McCraty & Shaffer, 2015; Thayer & Lane, 2000). A body of work focused on the central autonomic network (CAN) suggests that resting HRV reflects the brain’s ability to control the heart, other visceromotor organs, and neuroendocrine and behavioral responses that are critical for goal-directed behavior, executive function skills, adaptability, sustained health, and physiological self-regulation. Relatedly, increasing resting HRV through biofeedback was recently shown to induce memory bias for positive and neutral images, compared to negative (Cho et al., 2023). Conversely, they found that decreasing daytime HRV from baseline eliminated the positive and neutral biases observed in the increasing resting HRV condition, resulting in no memory biases across the positive, neutral, and negative memory stimuli. Interestingly, compared to positive memory stimuli, the neutral memory stimuli exhibited a numerically larger difference between the biofeedback conditions (increasing and decreasing resting HRV groups). Taken together with the current results demonstrating the HRV during REM was associated with greater neutral than negative memories, we suggest that the so-called “positivity bias” may be better characterized as a shift away from negative towards both positive and neutral memories. In fact, decreasing HRV and increasing spindle activity (with zolpidem) shifted memories towards the negative and away from neutral memories, suggesting a role for spindle activity in enhancing negative memories specifically. Further, including HRV during REM and sigma power in the regression model explained the most variance in the emotional memory tradeoff effect. Overall, our results indicate that increased HRV may have comparable roles during wakefulness and sleep in emotional memory processing. Specifically, to enhance the consolidation of neutral and positive memories, subtly shifting the focus away from negative memories.

REM is considered to be important for consolidating emotional memories (Goldstein & Walker, 2014; van der Helm & Walker, 2011; Walker, 2010). REM is thought to contribute to this process through greater connectivity between emotion regulation brain regions (i.e., amygdala and the PFC) during REM, compared to wake (van der Helm & Walker, 2011; Walker, 2010). Our results suggest that both neutral and emotional memory consolidation may occur simultaneously during REM, with a shift towards greater negative or neutral memories depending on the brain state, greater spindle activity versus greater HRV, respectively. Daytime HRV reflects how effectively an individual can activate the vagal nervous system to slow down the heart rate in response to increased sympathetic arousal from environmental stressors, or effectively transition from a heightened state to a more relaxed state (Friedman & Thayer, 1998; Thayer & Lane, 2000). Studies using biofeedback to increase HRV have demonstrated its causal role in improving emotion regulation by enhancing emotion regulation brain network connectivity (i.e., amygdala-PFC connectivity), thereby suggesting a link between top-down autonomic control and emotional regulation (Cho et al., 2023; Nashiro et al., 2023; Sakaki et al., 2016). These results support the notion that increasing HRV may enhance emotion regulation by improving feedback and communication between the CNS and ANS. This improved feedback may help interrupt negative thoughts or behavior and facilitate a shift towards emotion regulation, and thereby, a shift away from negative stimuli towards non-emotional and positive stimuli. Our results suggest that this process also occurs overnight through heightened vagal HRV and more effective emotion regulation during REM. Therefore, it appears that HRV during REM contributes to the emergence of a positivity bias and these processes are altered and diminished when vagal HRV is suppressed, as shown with zolpidem.

We propose that while REM is primarily responsible for emotional memory processing, vagal HRV during sleep plays a crucial role in shifting focus away from emotional memories, thereby facilitating the rescuing of non-emotional, neutral memories. One question that arises from this idea is: why would neutral memories be prioritized by HRV during sleep? Emotional memories are typically considered to be strong, more resistant to forgetting, and preferentially processed during sleep, compared to non-emotional, weaker memories (Payne & Kensinger, 2010; Wagner et al., 2001, 2006). It is possible that vagal HRV during sleep plays a crucial role in rescuing non-emotional, and comparatively weaker memories. Accordingly, encoding performance for neutral memories was negatively associated with HRV during sleep (both SWS and REM) in the placebo condition, and HRV during sleep then strengthened and integrated next day memory for these weak neutral stimuli. Prior work has similarly demonstrated a specific role of HRV during REM in enhancing the accessibility of weak memories and facilitating their incorporation into the existing network (Whitehurst et al., 2016; Yuksel et al., 2024). For example, Yuksel et al. found that, in participants with PTSD, disrupted REM and reduced REM HRV were linked to the failure to unlearn strong fear memories and replace them with weaker, extinction memories (Yuksel et al., 2024). These results similarly align with the model proposed by Norman et al., which posits that REM targets and strengthens weak memories, while NREM broadly enhances newly learned information (Norman et al., 2005). Based on these results, we posit that HRV during both SWS and REM may help facilitate the rescue of weak memories and then, HRV during REM further aids in refining these memories and integrating them into the existing network.

Our study has some limitations that should be considered when interpreting these results. First, we did not collect subjective valence or arousal ratings for the images, so we cannot assess changes in attributed emotional affect after sleep. This is particularly important because the majority of literature relates daytime HRV to emotional health, rather than emotional memory, as we explored here. Second, in our supplementary materials, we failed to replicate the wake findings from Cho et al. (2023) indicating higher resting state HRV leads to a positivity bias. These discrepancies may stem from a lack of positive images or varying experimental methods, such as the time of resting HRV recording (we recorded right before bed), and the ingestion of the drug right before our resting state, which could act as a confounding factor to influence HRV measurements, even if the pill was placebo. Therefore, future work should aim to disentangle the overnight processing of emotions and emotional memories with positive, negative, and neutral images. Third, we did not conduct an immediate memory test – the first test was administered after a delay of approximately 12 waking hours. There may have been changes in memory from initial learning to the first test that are not captured in this study. Fourth, this study did not have a habituation night, which may have led to the "first night effect" phenomenon, with increased disturbed sleep (Agnew et al., 1966). Although, to mitigate this effect, we counterbalanced picture sets, drug order, and randomized the presentation of image valences. However, we cannot fully discount its potential influence. Finally, although we did not directly measure respiration, we analyzed the high-frequency HRV peak to account for respiratory rate, which can impact HRV. HF peak showed no significant differences between the two drug conditions and varied within a narrow range (0.22 to 0.26 Hz), suggesting that respiratory activity is unlikely to have significantly influenced zolpidem’s effects on HRV and memory. Nonetheless, this possibility cannot be entirely ruled out.

## Conclusion

Our study provides evidence supporting a role of vagal HRV during REM in emotional memory processing. We show that higher REM HRV, not SWS HRV, significantly contributes to the overnight emergence of a positive memory bias, with greater memory for neutral images at the expense of negative images after a night of normal sleep. These results underscore the importance of considering physiological features into models of sleep-dependent memory consolidation. Additionally, given that higher daytime HRV has been linked to reductions in depression, anxiety, and stress, our findings highlight the importance of nighttime HRV in effective emotion regulation and its potential role in mitigating adverse mental health symptoms.

## Acknowledgments

We thank undergraduate research assistants in the laboratory for assistance with data collection. This work was supported by NIH Grant R01AG046646, Office 16 of Naval Research, and the Young Investigator Award to S.C.M. (Grant N00014-14-1-0513).

## References

Ackermann, S., & Rasch, B. (2014). Differential effects of non-REM and REM sleep on memory consolidation? Current Neurology and Neuroscience Reports, 14(2), 430. 10.1007/s11910-013-0430-8

Agnew, H. W., Webb, W. B., & Williams, R. L. (1966). The First Night Effect: An Eeg Studyof Sleep. Psychophysiology, 2(3), 263–266. 10.1111/j.1469-8986.1966.tb02650.x

Boardman, A., Schlindwein, F. S., Rocha, A. P., & Leite, A. (2002). A study on the optimum order of autoregressive models for heart rate variability. Physiological Measurement, 23(2), 325. 10.1088/0967-3334/23/2/308

Carr, M., & Nielsen, T. (2015). Morning REM Sleep Naps Facilitate Broad Access to Emotional Semantic Networks. Sleep, 38(3), 433–443. 10.5665/sleep.4504

Cellini, N., Torre, J., Stegagno, L., & Sarlo, M. (2016). Sleep before and after learning promotes the consolidation of both neutral and emotional information regardless of REM presence. Neurobiology of Learning and Memory, 133, 136–144. 10.1016/j.nlm.2016.06.015

Chen, P.-C., Niknazar, H., Alaynick, W. A., Whitehurst, L. N., & Mednick, S. C. (2021). Competitive dynamics underlie cognitive improvements during sleep. Proceedings of the National Academy of Sciences, 118(51), e2109339118. 10.1073/pnas.2109339118

Chen, P.-C., Sattari, N., Whitehurst, L. N., & Mednick, S. C. (2021). Age-related losses in cardiac autonomic activity during a daytime nap. Psychophysiology, 58(7), e13701. 10.1111/psyp.13701

Chen, P.-C., Whitehurst, L. N., Naji, M., & Mednick, S. C. (2020a). Autonomic Activity during a Daytime Nap Facilitates Working Memory Improvement. Journal of Cognitive Neuroscience, 32(10), 1963–1974. 10.1162/jocn_a_01588

Chen, P.-C., Whitehurst, L. N., Naji, M., & Mednick, S. C. (2020b). Autonomic/central coupling benefits working memory in healthy young adults. Neurobiology of Learning and Memory, 173, 107267. 10.1016/j.nlm.2020.107267

Cho, C., Yoo, H. J., Min, J., Nashiro, K., Thayer, J. F., Lehrer, P. M., & Mather, M. (2023). Changes in Medial Prefrontal Cortex Mediate Effects of Heart Rate Variability Biofeedback on Positive Emotional Memory Biases. Applied Psychophysiology and Biofeedback, 48(2), 135–147. 10.1007/s10484-023-09579-1

Cox, R., van Bronkhorst, M. L. V., Bayda, M., Gomillion, H., Cho, E., Parr, M. E., Manickas-Hill, O. P., Schapiro, A. C., & Stickgold, R. (2018). Sleep selectively stabilizes contextual aspects of negative memories. Scientific Reports, 8(1), Article 1. 10.1038/s41598-018-35999-9

Denis, D., Sanders, K. E. G., Kensinger, E. A., & Payne, J. D. (2022). Sleep preferentially consolidates negative aspects of human memory: Well-powered evidence from two large online experiments. Proceedings of the National Academy of Sciences, 119(44), e2202657119. 10.1073/pnas.2202657119

Electrophysiology Task Force of the European Society of Cardiology and the North American Society of Pacing and. (1996). Heart rate variability: Standards of measurement, physiological interpretation and clinical use. Circulation, 93, 1043–1065.

First, M., Gibbon, M., Spitzer, R., Benjamin, L., & Williams, J. (1997). Structured clinical interview for DSM-IV® axis ii personality disorders SCID-I. American Psychiatric Association Publishing.

Forte, G., Morelli, M., & Casagrande, M. (2021). Heart Rate Variability and Decision-Making: Autonomic Responses in Making Decisions. Brain Sciences, 11(2), Article 2. 10.3390/brainsci11020243

Friedman, B. H., & Thayer, J. F. (1998). Autonomic balance revisited: Panic anxiety and heart rate variability. Journal of Psychosomatic Research, 44(1), 133–151. 10.1016/S0022-3999(97)00202-X

Goldstein, A. N., & Walker, M. P. (2014). The Role of Sleep in Emotional Brain Function. Annual Review of Clinical Psychology, 10, 679–708. 10.1146/annurev-clinpsy-032813-153716

Groch, S., Wilhelm, I., Diekelmann, S., & Born, J. (2013). The role of REM sleep in the processing of emotional memories: Evidence from behavior and event-related potentials. Neurobiology of Learning and Memory, 99, 1–9. 10.1016/j.nlm.2012.10.006

Gujar, N., McDonald, S. A., Nishida, M., & Walker, M. P. (2011). A Role for REM Sleep in Recalibrating the Sensitivity of the Human Brain to Specific Emotions. Cerebral Cortex, 21(1), 115–123. 10.1093/cercor/bhq064

Hansen, A. L., Johnsen, B. H., & Thayer, J. F. (2003). Vagal influence on working memory and attention. International Journal of Psychophysiology, 48(3), 263–274. 10.1016/S0167-8760(03)00073-4

Jasper, H. (1958). Report of the committee on methods of clinical examination in electroencephalography. Electroencephalography and Clinical Neurophysiology, 10(2), 370–375. 10.1016/0013-4694(58)90053-1

Kaestner, E. J., Wixted, J. T., & Mednick, S. C. (2013). Pharmacologically increasing sleep spindles enhances recognition for negative and high-arousal memories. Journal of Cognitive Neuroscience, 25(10), 1597–1610. 10.1162/jocn_a_00433

Lang, P., J., Bradly, M., J., & Cuthbert, B., N. (1997). International affective picture system (IAPS): Technical manual and affective ratings (Vol. 1). NIMH Center for the Study of Emotion and Attention. https://scholar.google.com/scholar_lookup?&title=International%20affective%20picture%20system%20%28IAPS%29%3A%20Technical%20manual%20and%20affective%20ratings&journal=NIMH%20Center%20for%20the%20Study%20of%20Emotion%20and%20Attention&volume=1&pages=39-58&publication_year=1997&author=Lang%2CPJ&author=Bradley%2CMM&author=Cuthbert%2CBN

Lara-Carrasco, J., Nielsen, T. A., Solomonova, E., Levrier, K., & Popova, A. (2009). Overnight emotional adaptation to negative stimuli is altered by REM sleep deprivation and is correlated with intervening dream emotions. Journal of Sleep Research, 18(2), 178–187. 10.1111/j.1365-2869.2008.00709.x

Lipinska, G., Stuart, B., Thomas, K. G. F., Baldwin, D. S., & Bolinger, E. (2019). Preferential Consolidation of Emotional Memory During Sleep: A Meta-Analysis. Frontiers in Psychology, 10. 10.3389/fpsyg.2019.01014

Machado, C., Est’vez, M., P’rez-Nellar, J., Guti’rrez, J., Rodríguez, R., Carballo, M., Chinchilla, M., Machado, A., Portela, L., García-Roca, M. C., & Beltrán, C. (2011). Autonomic, EEG, and Behavioral Arousal Signs in a PVS Case After Zolpidem Intake. Canadian Journal of Neurological Sciences / Journal Canadien Des Sciences Neurologiques, 38(2), 341–344. 10.1017/S0317167100011562

Malik, M., & Camm, A. J. (1993). Components of heart rate variability—What they really mean and what we really measure. The American Journal of Cardiology, 72(11), 821–822. 10.1016/0002-9149(93)91070-X

McCraty, R., & Shaffer, F. (2015). Heart Rate Variability: New Perspectives on Physiological Mechanisms, Assessment of Self-regulatory Capacity, and Health risk. Global Advances in Health and Medicine, 4(1), 46–61. 10.7453/gahmj.2014.073

Mednick, S. C., McDevitt, E. A., Walsh, J. K., Wamsley, E., Paulus, M., Kanady, J. C., & Drummond, S. P. A. (2013). The Critical Role of Sleep Spindles in Hippocampal-Dependent Memory: A Pharmacology Study. The Journal of Neuroscience, 33(10), 4494– 4504. 10.1523/JNEUROSCI.3127-12.2013

Nashiro, K., Min, J., Yoo, H. J., Cho, C., Bachman, S. L., Dutt, S., Thayer, J. F., Lehrer, P. M., Feng, T., Mercer, N., Nasseri, P., Wang, D., Chang, C., Marmarelis, V. Z., Narayanan, S., Nation, D. A., & Mather, M. (2023). Increasing coordination and responsivity of emotion-related brain regions with a heart rate variability biofeedback randomized trial. Cognitive, Affective, & Behavioral Neuroscience, 23(1), 66–83. 10.3758/s13415-022-01032-w

Nishida, M., Pearsall, J., Buckner, R. L., & Walker, M. P. (2009). REM Sleep, Prefrontal Theta, and the Consolidation of Human Emotional Memory. Cerebral Cortex (New York, NY), 19(5), 1158–1166. 10.1093/cercor/bhn155

Norman, K. A., Newman, E. L., & Perotte, A. J. (2005). Methods for reducing interference in the Complementary Learning Systems model: Oscillating inhibition and autonomous memory rehearsal. Neural Networks, 18(9), 1212–1228. 10.1016/j.neunet.2005.08.010

Park, G., Van Bavel, J., Vasey, M., & Thayer, J. (2012). Cardiac Vagal Tone Predicts Inhibited Attention to Fearful Faces. Emotion (Washington, D.C.), 12. 10.1037/a0028528

Payne, J. D., & Kensinger, E. A. (2010). Sleep’s Role in the Consolidation of Emotional Episodic Memories. Current Directions in Psychological Science, 19(5), 290–295. 10.1177/0963721410383978

Payne, J. D., Kensinger, E. A., Wamsley, E., Spreng, R. N., Alger, S., Gibler, K., Schacter, D. L., & Stickgold, R. (2015). Napping and the Selective Consolidation of Negative Aspects of Scenes. Emotion (Washington, D.C.), 15(2), 176–186. 10.1037/a0038683

Payne, J. D., Stickgold, R., Swanberg, K., & Kensinger, E. A. (2008). Sleep Preferentially Enhances Memory for Emotional Components of Scenes. Psychological Science, 19(8), 781–788. 10.1111/j.1467-9280.2008.02157.x

Rechtschaffen, A., & Kales, A. (1968). A manual of standardized terminology, techniques and scoring system for sleep stages of human subjects. United States Government Printing Office.

Sakaki, M., Yoo, H. J., Nga, L., Lee, T.-H., Thayer, J. F., & Mather, M. (2016). Heart rate variability is associated with amygdala functional connectivity with MPFC across younger and older adults. NeuroImage, 139, 44–52. 10.1016/j.neuroimage.2016.05.076

Schäfer, S. K., Wirth, B. E., Staginnus, M., Becker, N., Michael, T., & Sopp, M. R. (2020). Sleep’s impact on emotional recognition memory: A meta-analysis of whole-night, nap, and REM sleep effects. Sleep Medicine Reviews, 51, 101280. 10.1016/j.smrv.2020.101280

Schumann, A., de la Cruz, F., Köhler, S., Brotte, L., & Bär, K.-J. (2021). The Influence of Heart Rate Variability Biofeedback on Cardiac Regulation and Functional Brain Connectivity. Frontiers in Neuroscience, 15. 10.3389/fnins.2021.691988

Simon, K. C., Whitehurst, L. N., Zhang, J., & Mednick, S. C. (2022). Zolpidem Maintains Memories for Negative Emotions Across a Night of Sleep. Affective Science, 3(2), 389–399. 10.1007/s42761-021-00079-1

Sinha, B., & Yadav, C. T. (2020). Effect of Zolpidem on sleep efficiency and heart rate during daytime nap. Indian Journal of Aerospace Medicine, 63(2), 83–89. 10.25259/IJASM_10_2019

Sotirchos, E. S., Fitzgerald, K. C., & Crainiceanu, C. M. (2019). Reporting of R2 Statistics for Mixed-Effects Regression Models. JAMA Neurology, 76(4), 507. 10.1001/jamaneurol.2018.4720

Suriya-Prakash, M., John-Preetham, G., & Sharma, R. (2015). Is heart rate variability related to cognitive performance in visuospatial working memory? [Preprint]. PeerJ PrePrints. 10.7287/peerj.preprints.1377v1

Thayer, J. F., Hansen, A. L., Saus-Rose, E., & Johnsen, B. H. (2009). Heart Rate Variability, Prefrontal Neural Function, and Cognitive Performance: The Neurovisceral Integration Perspective on Self-regulation, Adaptation, and Health. Annals of Behavioral Medicine, 37(2), 141–153. 10.1007/s12160-009-9101-z

Thayer, J. F., & Lane, R. D. (2000). A model of neurovisceral integration in emotion regulation and dysregulation. Journal of Affective Disorders, 61(3), 201–216. 10.1016/S0165-0327(00)00338-4

Trinder, J., Waloszek, J., Woods, M. J., & Jordan, A. S. (2012). Sleep and cardiovascular regulation. Pflügers Archiv - European Journal of Physiology, 463(1), 161–168. 10.1007/s00424-011-1041-3

van der Helm, E., & Walker, M. P. (2011). Sleep and Emotional Memory Processing. Sleep Medicine Clinics, 6(1), 31–43. 10.1016/j.jsmc.2010.12.010

Wagner, U., Gais, S., & Born, J. (2001). Emotional Memory Formation Is Enhanced across Sleep Intervals with High Amounts of Rapid Eye Movement Sleep. Learning & Memory, 8(2), 112–119. 10.1101/lm.36801

Wagner, U., Hallschmid, M., Rasch, B., & Born, J. (2006). Brief Sleep After Learning Keeps Emotional Memories Alive for Years. Biological Psychiatry, 60(7), 788–790. 10.1016/j.biopsych.2006.03.061

Walker, M. P. (2010). Sleep, memory and emotion. Progress in Brain Research, 185, 49–68. 10.1016/B978-0-444-53702-7.00004-X

Whitehurst, L. N., Cellini, N., McDevitt, E. A., Duggan, K. A., & Mednick, S. C. (2016). Autonomic activity during sleep predicts memory consolidation in humans. Proceedings of the National Academy of Sciences, 113(26), 7272–7277. 10.1073/pnas.1518202113

Whitehurst, L. N., Naji, M., & Mednick, S. C. (2018). Comparing the cardiac autonomic activity profile of daytime naps and nighttime sleep. Neurobiology of Sleep and Circadian Rhythms, 5, 52–57. 10.1016/j.nbscr.2018.03.001

Wiesner, C. D., Pulst, J., Krause, F., Elsner, M., Baving, L., Pedersen, A., Prehn-Kristensen, A., & Göder, R. (2015). The effect of selective REM-sleep deprivation on the consolidation and affective evaluation of emotional memories. Neurobiology of Learning and Memory, 122, 131–141. 10.1016/j.nlm.2015.02.008

Williams, P. G., Cribbet, M. R., Tinajero, R., Rau, H. K., Thayer, J. F., & Suchy, Y. (2019). The association between individual differences in executive functioning and resting high-frequency heart rate variability. Biological Psychology, 148, 107772. 10.1016/j.biopsycho.2019.107772

Yuksel, C., Watford, L., Muranaka, M., McCoy, E., Lax, H., Mendelsohn, A. K., Oliver, K. I., Daffre, C., Acosta, A., Vidrin, A., Martinez, U., Lasko, N., Orr, S., & Pace-Schott, E. F. (2024). REM disruption and REM Vagal Activity Predict Extinction Recall in Trauma-Exposed Individuals. bioRxiv, 2023.09.28.560007. 10.1101/2023.09.28.560007

Zhang, J., Yetton, B., Whitehurst, L. N., Naji, M., & Mednick, S. C. (2020). The effect of zolpidem on memory consolidation over a night of sleep. Sleep, 43(11), zsaa084. 10.1093/sleep/zsaa084

