## Supplementary Materials for "Vagal Heart Rate Variability During Rapid Eye Movement Sleep Reduces Negative Memory Bias"

#### Tables

| | Total (n = 33) | | | |
|---|---|---|---|---|
| | *%* | *n* | *M* | *SD* |
| Sex (female) | 48.5 | 16 | | |
| Age (years) | | | 20.5 | 2.92 |
| Ethnicity/Race | | | | |
| Hispanic or Latino | 39.4 | 13 | | |
| White | 33.3 | 11 | | |
| Asian | 21.2 | 7 | | |
| African American | 3 | 1 | | |
| Two or More Races | 3 | 1 | | |
| Education (years) | | | 14.5 | 2.29 |
| Health behavior/status | | | | |
| Caffeine/week | | | 0.91 | 1.16 |
| Alcohol/week | | | 0.73 | 1.18 |
| Habitual Napper | 60.6 | 20 | | |
| ESS | | | 6.36 | 2.69 |
| Weight | | | 155.4 | 29.2 |

**Table S1. Participant Characteristics at Baseline**

| Sleep | PBO | | ZOL | | |
|---|---|---|---|---|---|
| | Mean | SD | Mean | SD | p |
| TST | 537.948 | 48.003 | 540.450 | 39.194 | 0.785 |
| N2 (mins) | 283.000 | 53.399 | 288.950 | 46.487 | 0.441 |
| SWS (mins) | 110.362 | 39.192 | 121.866 | 39.834 | **0.039** |
| REM (mins) | 130.431 | 33.088 | 116.133 | 27.991 | **0.009** |
| WASO | 30.672 | 27.345 | 25.183 | 25.264 | 0.119 |

**Table S2. Descriptive Sleep Data**

| HRV | Stage | Measure | PBO | ZOL | p |
|---|---|---|---|---|---|
| HF (ln) | N2 | Mean (SD) | 6.568 (0.945) | 6.487 (0.935) | 0.545 |
| HF (ln) | SWS | Mean (SD) | 6.058 (1.105) | 6.060 (1.094) | **0.034** |
| HF (ln) | REM | Mean (SD) | 6.445 (1.011) | 6.446 (1.001) | 0.133 |
| RMSSD (ln) | N2 | Mean (SD) | 4.087 (0.542) | 3.970 (0.490) | 0.256 |
| RMSSD (ln) | SWS | Mean (SD) | 3.821(0.598) | 3.820 (0.593) | **0.024** |
| RMSSD (ln) | REM | Mean (SD) | 4.087 (0.542) | 3.970 (0.490) | 0.270 |
| Normalized HF | N2 | Mean (SD) | 0.534 (0.122) | 0.550 (0.135) | 0.533 |
| Normalized HF | SWS | Mean (SD) | 0.632 (0.155) | 0.630 (0.154) | **0.004** |
| Normalized HF | REM | Mean (SD) | 0.534 (0.122) | 0.550 (0.135) | 0.519 |

**Table S3. Summary of HRV Parameters Across Sleep Stages**

| | | | **Placebo** | | | **Zolpidem** | | |
|---|---|---|---|---|---|---|---|---|
| | | | **Mean** | **SD** | **d' (SD)** | **Mean** | **SD** | **d' (SD)** |
| Test 1 | Negative | Hit rate | 0.84 | 0.16 | 2.39 (.86) | 0.83 | 0.17 | 2.18 (.72) |
| | | False alarm | 0.17 | 0.17 | | 0.22 | 0.19 | |
| | Neutral | Hit rate | 0.76 | 0.18 | 2.05 (1.17) | 0.76 | 0.17 | 2.24 (1.17) |
| | | False alarm | 0.18 | 0.16 | | 0.14 | 0.17 | |
| Test 2 | Negative | Hit rate | 0.71 | 0.18 | 1.82 (.87) | 0.77 | 0.22 | 2.07 (1.01) |
| | | False alarm | 0.17 | 0.15 | | 0.19 | 0.20 | |
| | Neutral | Hit rate | 0.71 | 0.22 | 2.16 (.74) | 0.69 | 0.19 | 1.92 (.86) |
| | | False alarm | 0.11 | 0.12 | | 0.13 | 0.16 | |

#### Table S4. EPT Memory Performance

| | | | PBO | | | | ZOL | | | |
|---|---|---|---|---|---|---|---|---|---|---|
| EPT | HRV | Stage | r | p | t | CI | r | p | t | CI |
| Negative d' Test 1 | HF (ln) | REM | -0.240 | 0.244 | -1.197 | [-0.582, 0.169] | 0.130 | 0.524 | 0.646 | [-0.277, 0.502] |
| | HF (ln) | SWS | -0.260 | 0.218 | -1.268 | [-0.591, 0.155] | 0.070 | 0.741 | 0.334 | [-0.335, 0.452] |
| Neutral d' Test 1 | HF (ln) | REM | -0.600 | **0.001** | -3.522 | [-0.809, -0.260] | -0.032 | 0.876 | -0.158 | [-0.415, 0.359] |
| | HF (ln) | SWS | -0.570 | **0.004** | -3.161 | [-0.795, -0.204] | -0.096 | 0.654 | -0.455 | [-0.482, 0.319] |
| Negative d' Test 2 | HF (ln) | REM | -0.310 | 0.127 | -1.584 | [-0.631, 0.093] | 0.043 | 0.839 | 0.205 | [-0.359, 0.430] |
| | HF (ln) | SWS | -0.280 | 0.181 | -1.381 | [-0.606, 0.133] | -0.027 | 0.898 | -0.130 | [-0.418, 0.372] |
| Neutral d' Test 2 | HF (ln) | REM | -0.230 | 0.280 | -1.109 | [-0.580, 0.191] | 0.038 | 0.854 | 0.185 | [-0.355, 0.419] |
| | HF (ln) | SWS | -0.210 | 0.335 | -0.987 | [-0.573, 0.220] | 0.022 | 0.919 | 0.102 | [-0.385, 0.421] |

**Table S5. Correlations Between Vagal Activity and Emotional and Neutral Memory for Pre- and Post-Sleep**

| EPT | HRV | Stage | Drug | r | p | t | CI |
|---|---|---|---|---|---|---|---|
| Negative d' Test 1 | HF (ln) | Resting | PBO | -0.500 | 0.173 | -1.517 | [-0.874, 0.249] |
| Neutral d' Test 1 | HF (ln) | Resting | PBO | -0.390 | 0.333 | -1.054 | [-0.861, 0.429] |
| Negative d' Test 2 | HF (ln) | Resting | PBO | -0.340 | 0.365 | -0.970 | [-0.821, 0.414] |
| Neutral d' Test 2 | HF (ln) | Resting | PBO | -0.310 | 0.449 | -0.811 | [-0.835, 0.501] |
| Negative Overnight Performance Change | HF (ln) | Resting | PBO | 0.017 | 0.965 | 0.045 | [-0.655, 0.673] |
| Neutral Overnight Performance Change | HF (ln) | Resting | PBO | 0.250 | 0.555 | 0.624 | [-0.555, 0.810] |
| Emotional Tradeoff | HF (ln) | Resting | PBO | -0.150 | 0.676 | -0.433 | [-0.714, 0.528] |
| Negative d' Test 1 | HF (ln) | Resting | ZOL | -0.420 | 0.178 | -1.451 | [-0.800, 0.206] |
| Neutral d' Test 1 | HF (ln) | Resting | ZOL | -0.220 | 0.476 | -0.738 | [-0.686, 0.379] |
| Negative d' Test 2 | HF (ln) | Resting | ZOL | -0.370 | 0.236 | -1.262 | [-0.779, 0.258] |
| Neutral d' Test 2 | HF (ln) | Resting | ZOL | 0.028 | 0.928 | 0.092 | [-0.532, 0.570] |
| Negative Overnight Performance Change | HF (ln) | Resting | ZOL | -0.120 | 0.717 | -0.374 | [-0.648, 0.489] |
| Neutral Overnight Performance Change | HF (ln) | Resting | ZOL | 0.230 | 0.454 | 0.776 | [-0.370, 0.692] |
| Emotional Tradeoff | HF (ln) | Resting | ZOL | -0.340 | 0.208 | -1.324 | [-0.729, 0.203] |

**Table S6. Correlations Between Resting State Vagal Activity and Emotional and Neutral Memory**

### Figures

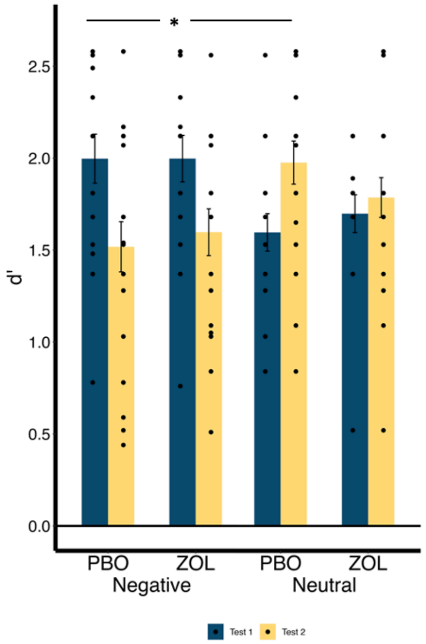


**Figure S1:** Memory performance (d') at test 1 and test 2 for placebo and zolpidem conditions. In the placebo condition, participants had a significant difference between test 1 and test 2 performance for negative images (p=.004), but not for neutral images (p=.928). In the zolpidem condition, there were no significant differences between test 1 and test 2 performance for negative or neutral images (p's>.132).

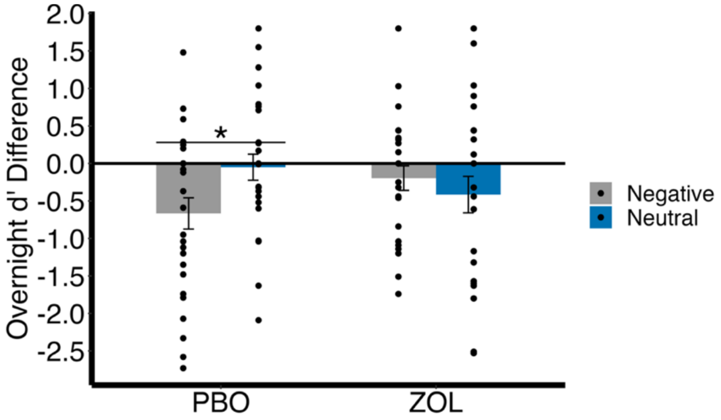


**Figure S2: Overnight EPT Memory Performance.** Participants had greater retention at test 1 than 2, as shown by a negative difference score. There was a significant three-way interaction between drug × test × emotion for the placebo condition, with greater memory for neutral than negative images (t=-2.463, p=.016, CI = [-1.146 -0.118]).

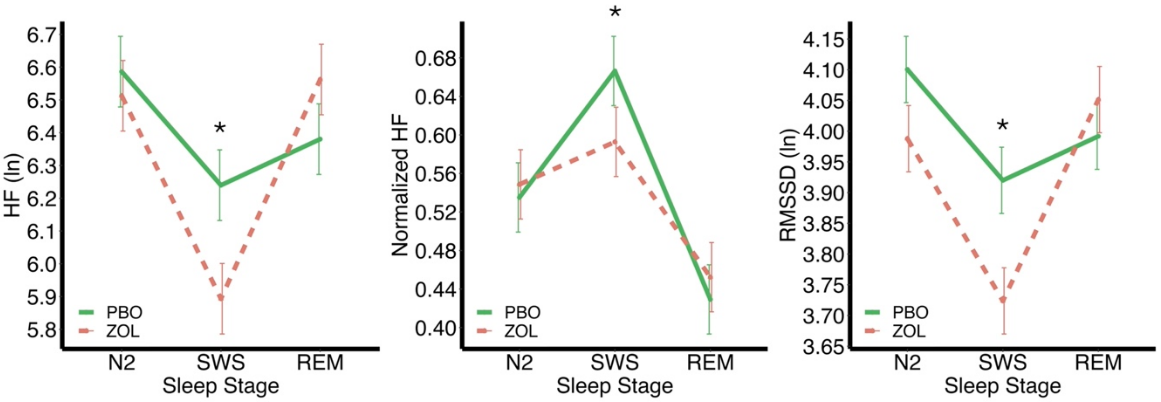


**Figure S3: Overnight HRV Modulated by Sleep Stage.** We report significant differences between drug conditions during SWS for (A) HF (ln) (p=.034), (B) Normalized HF (p=.004), and (C) RMSSD (p=.024). Yet, there were no significant differences between drug conditions during N2 or REM for (A) HF (ln) (p's>.133), (B) Normalized HF (p's>.519), and (C) RMSSD (p's>.256).

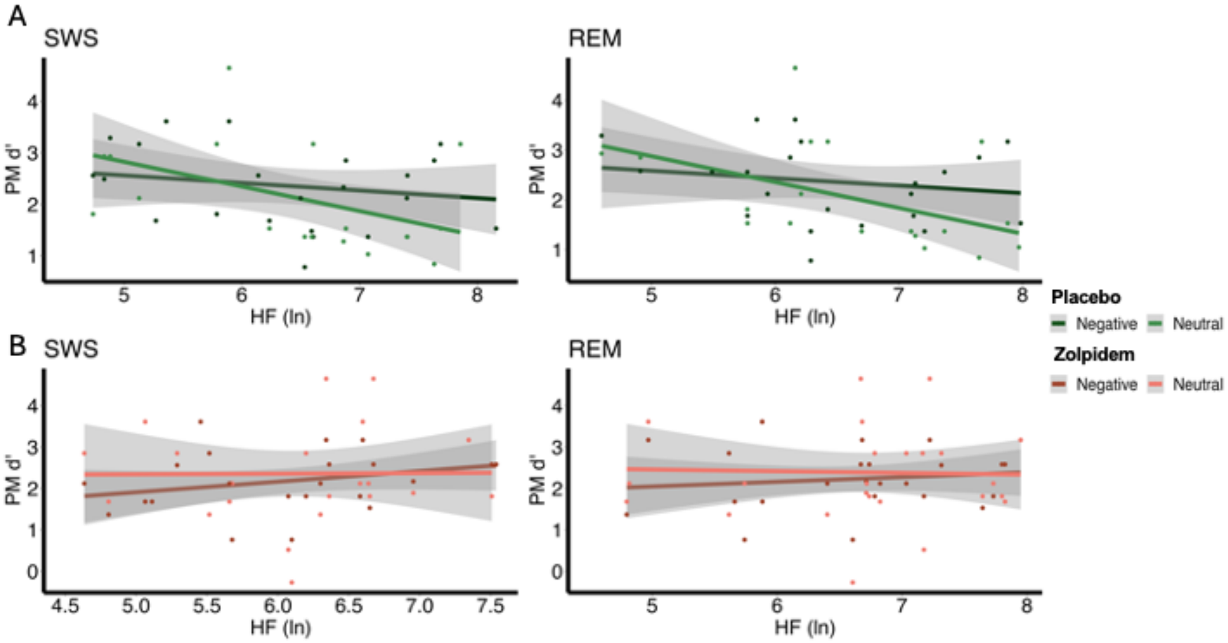


**Figure S4:** Correlations Between Vagal Activity and the Test 1 Performance (d') in Emotional and Neutral Memory. (A) In the placebo condition, pre-sleep test 1 memory performance for neutral images positively correlated with HRV during SWS (r= -0.570, p=.004) and with HRV during REM (r= -0.600, p=.001). However, no significant associations were found between negative images and HRV during SWS (r= -0.260, p=.218) or with HRV during REM (r= -0.0240, p=.244). (B) In the zolpidem condition, there were no significant correlations between test 1 memory performance for neutral or negative images and HRV during SWS (all p>.654) or with HRV during REM (all p>.524).

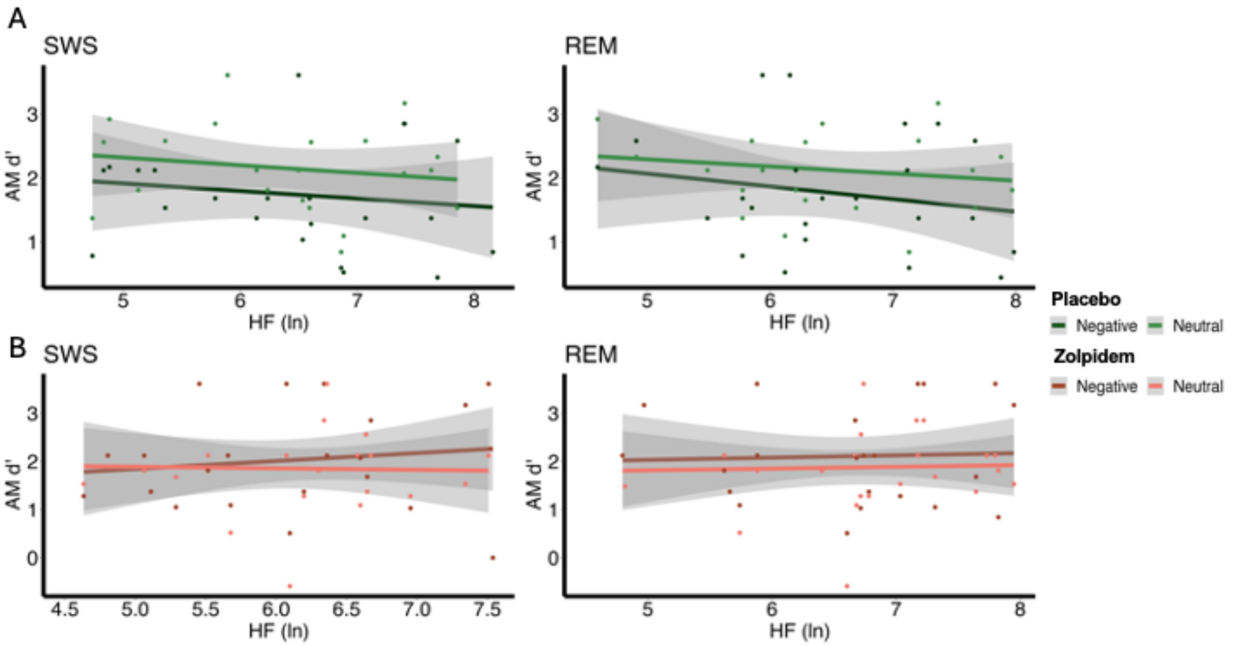


**Figure S5:** Correlations Between Vagal Activity and the Test 2 Performance (d') in Emotional and Neutral Memory. (A) In the placebo and (B) zolpidem conditions, there were no significant correlations between test 2 memory performance for neutral or negative images and HRV during SWS (all $p>.181$) or with HRV during REM (all $p>.127$).
